# Assigning credit where it’s due: An information content score to capture the clinical value of Multiplexed Assays of Variant Effect

**DOI:** 10.1101/2023.10.20.562794

**Authors:** John Michael O Ranola, Carrie Horton, Tina Pesaran, Shawn Fayer, Lea M. Starita, Brian H Shirts

**Author notes:** (Corresponding Author) Brian Shirts. Author Emails: John Michael Ranola – Carrie Horton – Tina Pesaran – Shawn Fayer - Lea Starita –.

## Abstract

**Background:** A variant can be pathogenic or benign with relation to a human disease. Current classification categories from benign to pathogenic reflect a probabilistic summary of current understanding. A primary metric of clinical utility for multiplexed assays of variant effect (MAVE) is the number of variants that can be reclassified from uncertain significance (VUS). However, we hypothesized that this measure of utility underrepresents the information gained from MAVEs and that an information theory approach which includes data that does not reclassify variants will better reflect true information gain. We used this information theory approach to evaluate the information gain, in bits, for MAVEs of *BRCA1, PTEN*, and *TP53*. Here, one bit represents the amount of information required to completely classify a single variant starting from no information.

**Results:** *BRCA1* MAVEs produced a total of 831.2 bits of information, 6.58% of the total missense information in *BRCA1* and a 22-fold increase over the information that only contributed to VUS reclassification. *PTEN* MAVEs produced 2059.6 bits of information which represents 32.8% of the total missense information in *PTEN* and an 85-fold increase over the information that contributed to VUS reclassification. *TP53* MAVEs produced 277.8 bits of information which represents 6.22% of the total missense information in *TP53* and a 3.5-fold increase over the information that contributed to VUS reclassification.

**Conclusions:** An information content approach will more accurately portray information gained through MAVE mapping efforts than counting the number of variants reclassified. This information content approach may also help define the impact of modifying information definitions used to classify many variants, such as guideline rule changes.

## Background

Variant classification guidelines for human disease typically leads to a binary outcome. A variant can either be pathogenic or benign for a human disease. While there are cases where this pathogenic/benign dichotomy may not be suitable, it is appropriate for many genes with clear disease or risk phenotypes. The familiar 5 categories of pathogenic, likely pathogenic, uncertain/VUS, likely benign, and benign attempts to add a measure of certainty of the outcome to the pathogenic/benign dichotomy(1). Rather than putting all the variants that are thought to be pathogenic in a single category, they are divided into ones with higher and lower probability to be pathogenic, called pathogenic and likely pathogenic, respectively. The same applies for benign and likely benign variants.

This type of categorization that considers the certainty of the classification bears a striking resemblance to information theory, where a key measure is the quantification of uncertainty, which is called entropy. By borrowing ideas from information theory, these 5 categories can be replaced by a continuous value, information content, which can precisely convey the certainty of the classification of variants. Specifically, we can convert the probability of pathogenicity to a measure of information content(2,3). This information content, which ranges between 0 and 1, is often measured in bits. In this context, a bit with a value of 0 means that there is no certainty in the classification of the variant (i.e. it has a 50-50 chance of being pathogenic or benign, alternatively it has a probability of pathogenicity of 0.5). On the other hand, a bit with a value of 1 means that we are completely certain of the classification of the variant (i.e. the variant is 100% pathogenic or 100% benign, alternatively it has a probability of pathogenicity of 0 or 1). It would mean that there is complete evidence for classification either toward pathogenicity or benignity.

This shift from discrete categories to a continuous value adds both mathematical and practical benefits. For example, the information content yield of multiplexed assays of variant effect (MAVE) can be precisely quantified. In a MAVE, a great number of variants are measured for functional effects, however the clinical value of these studies is usually not measured in the number of variant effects measured, but the number of variants reclassified from VUS to a more certain category. In these studies many variants do not shift classification categories, thus a large amount of the information gained is not counted (4,5) (Figure 1 modified from Vollmer 2007(2)). An information content framework allows one to quantify information for both variants that are reclassified and variants that are not. In this way, the full information content of the study can be assessed. Full information content of a MAVE, or MAVE information score, can be determined by summing up the changes in certainty of all variants in the study. Change in information depends on what information is already present and what is added. If a variant is already clearly classified, adding more evidence does not meaningfully increase the information about that variant. Similarly, the information content framework incorporates evidence that goes against prior evidence. New evidence supporting pathogenicity for a likely benign variant decreases the classification certainty and counts as negative information. Incorporating all evidence is important for ultimate accuracy, even if it may temporarily decrease the certainty of classification until more evidence arises. These situations may not be reported in current literature, particularly when there is no shift from one classification to another; using an information framework encourages full reporting of the effects of all evidence on probability of pathogenicity.

**Figure 1.**
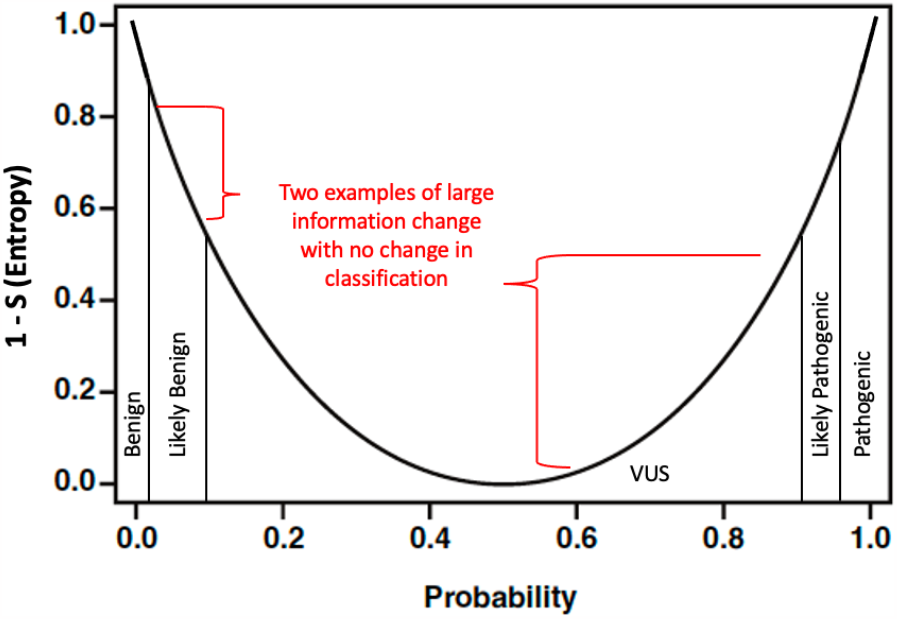
Conversion of probability of pathogenicity to information content modified from Vollmer 2007(2)

One feature of this method is that the resulting value is easily interpretable and compared across studies since a value of 1 bit represents 1 variant moved from complete uncertainty (50% chance of either benign or pathogenic) to complete certainty. That means that if the calculated information content of a study is 30 bits, then it is equivalent to completely classifying 30 variants from being completely uncertain. Note that depending on the study, the same number of bits of information content could be the result of gaining a small amount of information from many variants (as is common for MAVEs,) or a large amount of information from a of a small number of variants.

MAVE information score can be further incorporated into a gene-wide view. A gene can be thought of as a string of information where we know all the information for the reference strand since all positions in the reference should be benign. We don’t know all information for all possible variant strings. If each possible variant represents a bit of information, then the number of possible variants in a gene is the number of bits a gene needs to be completely classified. One can define this number as the finish line for completely classifying all possible single nucleotide variants in a gene. If a MAVE’s total information content is divided by this amount, the percentage of all possible information gained is calculated for that study. Alternatively, one can sum the bits for every variant in the gene from a MAVE and all other evidence sources to show overall progress for variant interpretation within that gene.

In this study, we explain the principles of applying information content to MAVEs and illustrate these principles by calculating the proportion of total missense variant information in a gene that several functional studies provide. We demonstrate that information content can capture much more than just changes in classification. We also illustrate how information content calculations can quantify the effect of changing prior probability or altering variant classification guidelines on apparent gene-wide information. The information framework can also be used to prioritize genes for MAVEs.

## Results

### Application 1 Quantifying the total information content of MAVEs

Total information content generated was calculated for several MAVEs that assessed variants in *BRCA1, PTEN*, and *TP53*(6). We used date from prior analysis of MAVEs and prior proposed translation of MAVE data to ACMG rules (see methods). Evidence criteria were converted to odds path per Tavtigian et. al.(7) then to posterior probability using standard Bayesian calculation. Posterior probability was converted to information content (see methods).

We compared information gain while only considering changes that resulted in VUS reclassification to the information gain from all single nucleotide substitutions reported in MAVE data. Some MAVEs evaluate amino acid changes that require more than one DNA substitution, although such data can be used in the proposed framework, we exclude it for this paper.

The *BRCA1* MAVE (8) examined 3893 variants with 2821 functional, 249 intermediate, and 823 showing loss of function. Functionally normal classification was considered strong benign evidence and loss of function was considered strong pathogenic evidence. Conversion of this evidence to posterior probability using prior odds of 0.1 and then to information content resulted in 813.2 bits of information gained by the study.

The *PTEN* MAVEs (9,10) examined 8198 variants for effects on protein abundance and examined 7657 variants for activity. The number of overlapping variants was 7639, of which 4811 had combined scores which were considered strong pathogenic evidence and 303 which were considered benign supporting evidence. Conversion of this evidence to posterior probability using prior odds of 0.1 and then to information content yielded a total information content of 893.6 bits of variant classification information added by the study for changes possible through single missense substitutions.

Four *TP53* MAVEs were combined and used to train a naïve bayes classifier and make predictions on 7893 variants with scores in each of the four assays(6,11,12). The Bayes classifier predicted 5070 as normal and 2823 as abnormal. These were assigned weights of benign moderate and pathogenic strong evidence, respectively. Conversion of this evidence to posterior probability using prior odds of 0.1 and then to information content yielded 160.0 bits of variant classification information for changes possible through single missense substitutions.

Comparing limited reports that include only VUS reclassified to our method which includes all classification information generated by several MAVEs, the information content increased 22-fold for the *BRCA1* assay, 85-fold for the *PTEN* assays, and 3.5-fold for *TP53* assays. (See Table 1)

**Table 1:** Information content in bits for reclassifying VUS as presented in Fayer et al.(6), total information content from all single substitutions reported with functional data presented in papers originally listing data, and total possible missense information.

|  | VUS only<br>(Fayer et<br>al.) | Single<br>substitution | Total bits of<br>missense<br>information in<br>gene |
| --- | --- | --- | --- |
| *BRCA1* | 36.9 | 813.2 | 12,351 |
| *PTEN* | 10.5 | 893.6 | 2,721 |
| *TP53* | 46.3 | 160 | 2,571 |

### Application 2 Quantifying the proportion of total missense information content of a gene generated by a MAVE

Total missense information content of a gene can be calculated by counting the number of possible missense variants in a gene. Since each variant represents one bit of information, the total missense information content of a gene, in bits, is the number of possible missense variants. These were 12,351; 2,721; and 2,571 for *BRCA1, PTEN*, and *TP53* respectively. Using the information generated by MAVEs for these genes listed above, the MAVEs generated 6.7%, 32.8%, and 6.2% of total possible single-substitution variant classification information, for *BRCA1, PTEN*, and *TP53* respectively. (See Figure 2)

**Figure 2.**
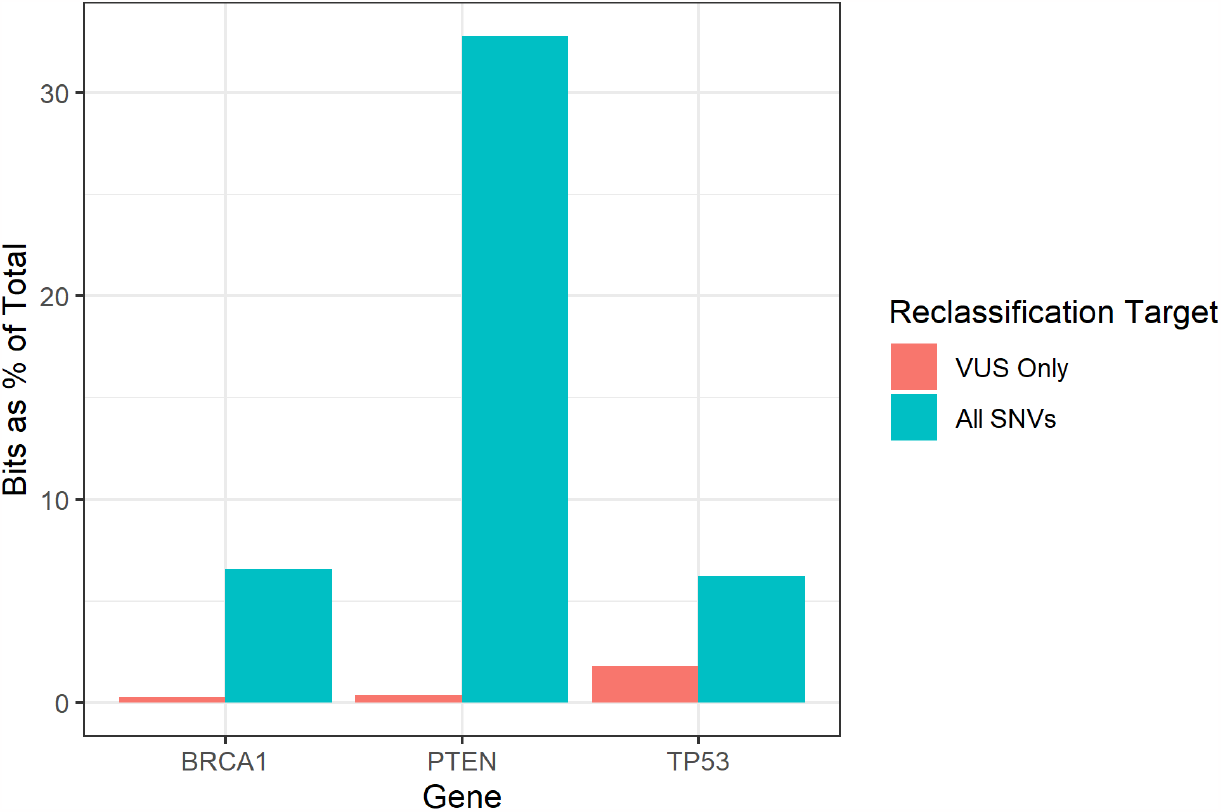
Percent of total information found by each study. A plot of the percent of total information found by each of the studies.

### Application 3 Quantifying the apparent information effect of a classification guideline rule change

For well-established functional studies 2015 ACMG-AMP guidelines recommend using evidence codes PS3 and BS3 indicating strong evidence for or against pathogenicity, respectively(13). However, different levels of evidence have been proposed for MAVEs that meet stronger or weaker validation criteria(4,14–17). If variant classification guidelines or strength of evidence criteria change, the changes have an effect on the apparent information for any variant that is impacted by the guideline change. For example, evidence against pathogenicity for *PTEN* MAVEs is considered supporting evidence primarily because there are very few established benign *PTEN* variants to use in validation. If validation data grows the evidence level for the MAVE may change. This would result in an apparent change in information content for many variants. Similarly, if ACMG-AMP committees or ClinGen VCEPs decide to refine the level of evidence assigned to a specific rule, there are many variants for which the apparent information content would change. We evaluated how applying single ACMG-AMP evidence levels combined with different priors using the Tavtigian et al.(7) Bayes framework would result in different levels of evidence (Figure 3). This analysis illustrates how the greatest information gains always occur when the prior probability is 0.5. Differences between points on the same vertical prior line show how shifting evidence assignment will change apparent information. This also illustrates that applying benign evidence to a variant with a high prior probability results in an apparent information loss from the increase in uncertainty or increasing entropy (all points with information change below 0 as plotted on y-axis). There is a similar result when pathogenic evidence is applied to a variant with a low prior probability.

Figure 3 plots how different evidence levels combined with prior probabilities result in different amounts of apparent information.

**Figure 3.**
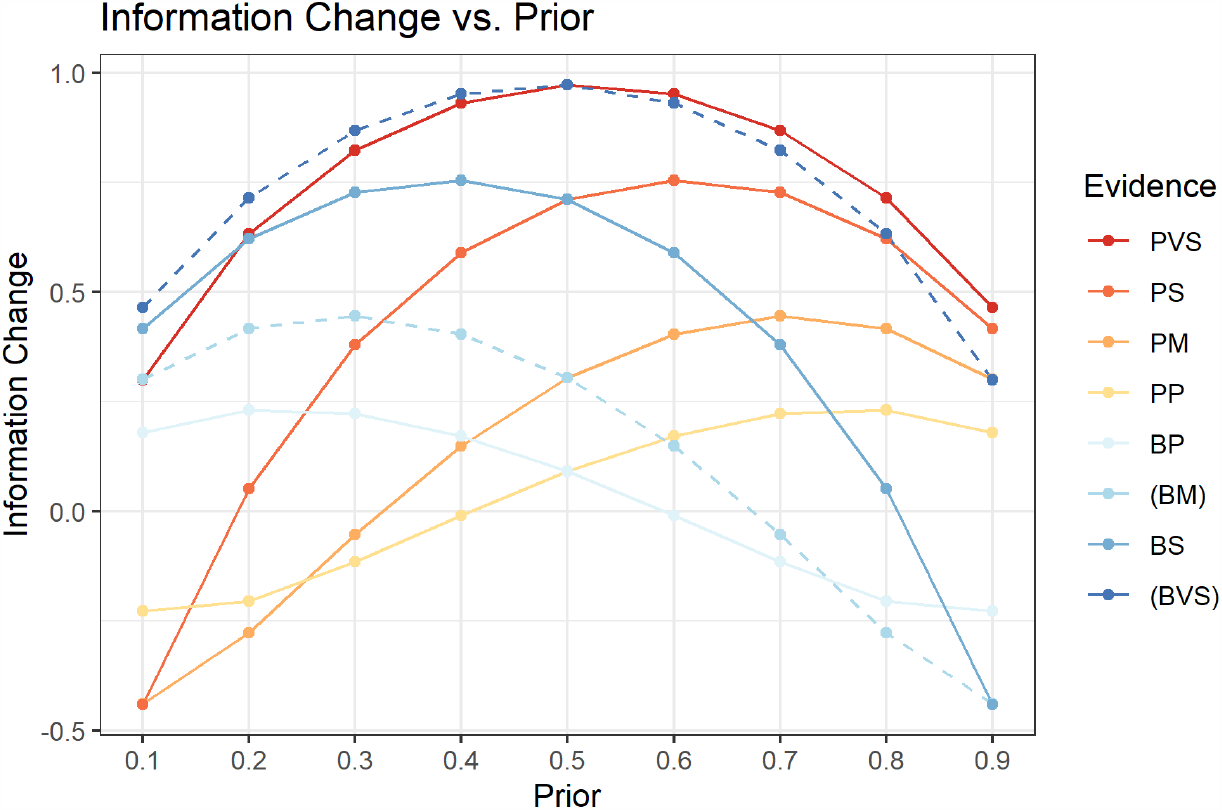
Plot of the information change for the different types of evidence across different prior probabilities. Information content loss for pathogenic evidence occurs at lower prior probabilities since these priors contain high information content for benign interpretation. Incorporation of the pathogenic evidence for a variant with a low prior moves the probability in the pathogenic direction and towards greater uncertainty. The same effect occurs for benign evidence with a high prior probability of pathogenicity since incorporation of benign evidence will reduce the probability towards a more uncertain class, thus reducing information content. The listed evidence categories are pathogenic very strong(PVS), pathogenic strong(PS), pathogenic moderate(PM), pathogenic supporting(PP), benign supporting(BP), benign moderate(BM), benign strong(BS), and benign very strong(BVS). Note that benign moderate and benign strong are not currently approved categories but are listed in parentheses for completion.

## Discussion

Currently, the clinical utility of MAVEs is measured as the number of individual variants classified or reclassified from variant of uncertain significance (VUS) to something else. Under current practice, a study which reclassifies a small number of variants may be perceived as more useful than a study which provides evidence for many variants but not enough for any one variant to be reclassified. An information content framework for evaluating MAVEs will provide a more accurate measurement of the full contribution of the functional data evidence. We have shown that reporting only changes in classification under-values the information yield of a MAVE, often by more than an order of magnitude.

In this study we focused on the change in relative information content, but absolute information content of a study can be quantified by calculating information content using a prior of 0.5 which assumes no prior information. This effectively eliminates any conflicting information and thus only yields positive information. With relative information content a study can seemingly provide negative information for individual variants due to the new evidence conflicting with the prior evidence (See Figure 3). While the idea of negative information content may seem odd at first glance, intuitively it makes sense that new conflicting information leads to a loss of certainty. It can also be thought of as the new information cancelling out previous information. This is evident in the *TP53* analysis where variants yielded 0.3 bits if they were functionally normal and -0.44 bits if they were functionally abnormal assuming a 0.1 prior. Since there were 1181 functionally abnormal variants, this led to a low total information content for the study. If we instead assumed a prior of 0.5, the variants would yield 0.3 bits if they were functionally normal and 0.71 for functionally abnormal. This would result in a total of 1518.61 bits of information instead of the 160 bits listed in Table 1.

In addition, an information content framework allows gene-level reporting of information in that it enables quantification of the total percentage of information provided by a MAVE for a given gene. The ability to quantify information as a percentage of a whole on a gene level has several benefits: it defines the variant information that exists, it can effectively illustrate the proportion of total information a specific MAVE has produced, and it can accurately illustrate what proportion of information remains missing for a gene or a variant. In the context of relative information, it can help prioritize future MAVEs by showing genes where a small proportion of total information exists and where efforts are likely to lead to a large increase in relative information.

Finally, understanding how classification rules influence apparent information will help committees that arbitrate guidelines for variant classification to quantify the effect of changing any specific rule more accurately across many variants in a gene. In addition to scrutinizing the effect on a few variants, summing or averaging the change of information over all variants can give a global assessment of the information change implied by the change in the strength assigned to evidence rules. The goal of classification guidelines should be to accurately quantify the information from different sources. Knowing how specific individual variants are classified is a clinical imperative; however, using a variant information framework shows how high-throughput methods and rule modifications can change the entire information landscape of the gene. Evaluating changes in information content with a classification rule change across many variants might help committees that propose changes to guidelines determine if changes appropriately reflect expected information content.

## Conclusions

Counting the number of VUS reclassified is a dramatic underestimate of the information content of MAVEs. Variant reclassification information reported as a change in information content provides a more accurate representation of the true information gained. We believe that incorporating an information content framework when presenting reclassification evidence will lead to a better understanding of the value of different classification projects and, in the end, better classification of variants. It will also allow quantification of how much information is missing for different genes allowing prioritization of genes with less relative knowledge. Reporting information content will better allow geneticists to understand the contributions of high-throughput information sources and guideline rule changes more wholistically. Better classification in turn leads to better clinical validity and utility of genetic testing and genetic risk assessment.

## Methods

### Theoretical background

Each specific variant in a gene is either pathogenic or benign for the disease associated with the gene. Each missense variant is assumed to be either pathogenic or benign and thus contain on bit of pathogenicity information. For simplicity we will consider that each gene of interest has a single defined gene-disease relationship. Variant classification can be described as gathering information about binary choice. For simplicity we also ignore the issues of reduced penetrance, as current classification systems have also done. Probability of pathogenicity can be converted to information entropy and then to a measure of information content(2).

Entropy, S, is defined in the equation below where p is the probability of pathogenicity.

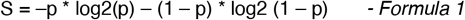

Information content is defined as 1-S or equivalently the difference between Smax and S.

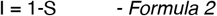

Information content provides a measure of certainty of outcome rather than the outcome itself. If there is uncertainty about any bit of information the amount of uncertainty can be expressed as a fraction of a bit of information being available. In this case, a 50% probability of pathogenicity amounts to 0 bits of information. On the other hand, either 0% or 100% probability of pathogenicity amounts to 1 bit of information (See Figure 1).

Conceptually, thinking of variant classification within the entropy framework as bits of information allows new mathematical transformations. For example, bits of information can be summed and averaged across a larger message or gene. We describe several examples of these applications below.

#### Application 1 Information Content of MAVEs

Total information content generated by several MAVEs was calculated. These assessed variants in the genes *BRCA1, PTEN*, and *TP53*. Information content gain depends on the information already available from current information, including the prior. For the VUS in Fayer et al.(6), the information content of the MAVEs were calculated as the change in information content for the specific variant with and without the evidence that the MAVE provided. We used date from prior analysis of MAVEs and prior proposed translation of MAVE data to ACMG rules. Data for *BRCA1* was acquired from Findlay et al. 2018(8) (Supplementary file 2). Data for *PTEN* was acquired from Matreyek et al. 2018(9) (online: http://abundance.gs.washington.edu) and Mighell et al. 2018(10) Table S6. Data for *TP53* was acquired from Table S4 from Fayer et al. 2021(6).

Briefly, MAVE evidence was assigned to ACMG classification criteria as described previously and then converted to odds path as described in Tavtigian et al.(7) and implemented in Fayer et al.(6) Odds Path were combined with prior probability to yield posterior probability of pathogenicity. This posterior probability of pathogenicity for each variant was converted to entropy and information content using formulas 1 and 2 (see Supplemental Tables S1, S2, S3 in Fayer et al) (6). For the *PTEN* and *TP53* MAVEs, information content was calculated using a prior pathogenicity of 0.1 as suggested in Tavtigian et al(7). Protein variants reported by other MAVE assays were subset into those that can be achieved through a single substitution and those that require more than one DNA change.

#### Application 2 Total Missense Variant Information Content in a Gene

To calculate the total information content of all missense variants in a gene, we took amino acid sequences for the *BRCA1, PTEN*, and *TP53* from the first entry in the gene’s respective UniProt(18). We then looked at each amino acid, found the RNA codons that could code for it, and counted the average number of possible missense, nonsense, and synonymous changes that could be achieved through a single nucleotide substitution. We then used amino acid sequence data for each gene from UniProt (18) and counted the number of possible missense variants for the gene which is equivalent to the total information content of a gene. Some *PTEN* and *TP53* MAVEs include assessment of amino acid changes that require more than one missense variant. Those assays define a different information space, which is larger than the single missense information space considered in this manuscript.

#### Application 3 Quantifying the apparent information effect of a classification guideline rule change

To evaluate how classification guideline rule changes might alter the apparent information gained from applying those rules, we applied each rule across the range of possible prior probabilities and plotted the information content from applying the single rule strength to the prior.

## Acknowledgements

The work was funded by National Institutes of Health’s National Human Genome Research Institute grants 1R01HG013025 to BHS and LMS and 5UM1HG011969 to SF and LMS.

